# The Coevolutionary Romance of Social Learning and Parasitic Behavior

**DOI:** 10.1101/055889

**Authors:** Richard McElreath

## Abstract

Once an animal begins to acquire behavior by social learning, it may be seduced by parasitic parasitic, behavior that reduces the animal’s fitness and thereby increases its own spread. However, the animal’s psychology will coevolve, potentially limiting the influence and spread of parasitic behavior. I revisit prominent models of the evolution of social learning and introduce the possibility of parasitic behavior. First, I explore a courtship between primitive social learning and parasitic behavior. Parasitic behavior can spread, but selection on the host then reduces social learning and limits its importance. Both parties are frustrated. In the second part, I study a reconciliation dynamic in which social learning becomes strategic about who it partners with. In this model, parasitic behavior can become prevalent and substantially reduce host fitness. However, it may also evolve to be mutualistic and raise the mean fitness of the host organism. When this occurs, natural selection may favor psychological susceptibility to parasitic behavior. Both social learning and socially learned behavior can enjoy a happy ending.

## 1. INTRODUCTION

A recurring idea about the evolution of behavior, especially human behavior, is that behavior itself may be subject to natural selection. Since humans acquire much behavior through social transmission from people other than their parents and close blood relations, behavior that manipulates human psychology may spread, even if such behavior imposes a fitness cost on those who perform it (Dawkins 1976, Cavalli-Sforza and Feldman 1981, Boyd and Richerson 1985, Durham 1991). Some behavior may fit human cognition in ways that make harder to forget and easier to transmit. If a variant of such behavior appears that can harness human energy for its own reproduction, such costly behavior may spread at the expense of neutral or adaptive behavior. Call such behavior *parasitic*, indicating that these variants manipulate human learners in ways that impose costs but also favor their own spread. Possible examples include dangerous hobbies such as rock climbing and drug abuse, professional occupations that require postponing reproduction, and religious practices and superstitions that encourage self-sacrifice.

Why can’t human psychology reliably reject such behavior? Richerson and Boyd (2005, Chapter 5) suggest an analogy with breathing: Despite many millions of years for evolution to perfect breathing, every breath still exposes us to countless potential pathogens. The lungs are defended, but remain penetrable, because breathing requires some permeability. Social learning entails a similar dilemma: Learning from others requires some credulity, and so our psychology may be defended, but a clever idea can nevertheless penetrate it.

This argument does not suggest that brains are helpless victims of parasitic behavior. Any psychology capable of sophisticated social learning must have coevolved in the shadow of such parasitic behavior. And so the design of social learning, whether innate or rather developmentally acquired, reflects a tradeoff between the acquisition of adaptive behavior and defense against manipulation. Human social learning is strategically rich and powerfully inferential (Boyd and Richerson 2000). Methods of learning transform both the contents and frequencies of behavior, and behavior transforms methods of learning. This sets up an essentially game-theoretic interaction that is difficult to reason through.

This paper does not attempt to shed any new light on psychological mechanisms or empirical evidence for parasitic behavior. Nor does it present a complete framework for explaining behavioral dynamics. Instead, it uses dynamic models to modestly address the logic of the core parasitic behavior hypothesis: Natural selection on behavior may favor parasitic variants that impose fitness costs but nevertheless spread; at the same time, the psychology that makes social learning possible should coevolve under the threat of parasitic behavior. What are the consequences of this coevolutionary process?

I address these questions by revisiting prominent models of the evolution of social learning and introducing the possibility of parasitic behavior. Few population dynamic models exist in which natural selection on behavior itself can spread costly behavior (Kendal and Laland 2000). The literature on gene-culture evolution for its part has tended to focus instead on adaptive learning mechanisms and understanding cumulative cultural evolution (Henrich and McElreath 2003). Boyd and Richerson (1985) do present an illustrative parent-teacher model that focuses on the conflict between genetic and cultural fitness (Chapter 6) and a model of runaway status signaling (Chapter 8). But the social learning systems in those models do not coevolve with behavior.

The new models I present here attempt to be the simplest that allow coevolution between parasitic behavior and social learning. While simple, these models are still more detailed than the most elaborate verbal models. Even ignoring many of the non-selective forces that matter in real cultural systems, I demonstrate that parasitic behavior and social learning may coevolve in surprising ways, owing to the strategic nature of some forms of social learning. The coevolution of parasitic behavior and social learning is analogous to a romance, in which behavior wants the learner to fall in love, but the easiest path to the learner’s heart may transform the behavior and have unanticipated consequences for both parties. Either or both behavior and psychology may wind up frustrated or rather elated.

I present the models in two parts: (1) an initial “courtship” phase in which a psychology for simple social learning meets parasitic behavior for the first time and (2) a “reconciliation” phase in which social learning becomes more choosy about which behavior it partners with. In the courtship phase, I insert the simplest formulation of parasitic behavior into the simplest and best-studied model of the evolution of social learning. This model is deeply unsatisfactory, but it is a necessary step, and it proves valuable. Parasitic behavior can spread in this model, but never very far. The presence of parasitic behavior leads the population to a reduced reliance on social learning. In the reconciliation phase, social learning becomes choosy about which behavioral variants it adopts. Social learning is deployed strategically rather than blindly. Social learning is much more adaptive in this model, and parasitic behavior can also spread much further, without reducing reliance on social learning. But parasitic behavior may also become positively associated with adaptive behavior and actually allow the population to reach higher levels of adaptation than in the absence of parasitic behavior.

If you are not thrilled by the notion of reading mathematical models, I sympathize. I have confined most of the mathematics in an appendix—where they rage with logic for your later inspection—and instead illustrated the main text with colorful simulation examples. A great deal can be gathered by reading the model assumptions and figure captions, before investing in deep perusal.

## 2. SIMPLE SOCIAL LEARNING AND PARASITIC BEHAVIOR

To say that natural selection has favored *social learning* means only that selection has favored heritable traits that lead individual behavior to be influenced by the behavior of other individuals. This influence may take diverse forms and have diverse consequences (Henrich and McElreath 2003, Laland 2004). *Simple* social learning means herein a form of social influence in which an individual acquires a behavioral variant in proportion to its frequency in the population of social influencers. Simple social learning saves on the costs of exploring the environment but risks using inappropriate behavior for the current environment. It evolves when the costs of innovation are relative high and the environment is not too variable, either in space or time (Richerson and Boyd 2005). The simplest model that justifies these inferences is a 1988 model by Alan R. Rogers (Rogers 1988). This model has been the basis of much discussion and many elaborations, because it is one of the simplest models that satisfactorily represents the basic dilemma of social learning. It is a good example of an argument that lacks nuance but has had tremendous positive impact.

The model I present in this section diverges very little from the Rogers model. This has two advantages. First, the model I present here is among the simplest models of the evolution of parasitic behavior that also allows the coevolution of social learning. Second, the model retains comparability not only to Rogers’ original model, but also to the large literature that relates to it.

### 2.1. Model description

Suppose a large haploid population of organisms with a simple life cycle: They are born, have the opportunity to learn about the current environment, use what they’ve learned to acquire resources, use those resources to reproduce, and finally remain alive just long enough for the next generation to observe their pattern of behavior. Learning can be either *individual* (I) or *social* (S) and is coded for by a single locus with two alleles. Individual learners pay a fitness cost *c* for a chance *s* of acquiring behavior that is adapted to the current state of the environment, earning them *b* units of fitness. Social learners instead acquire the behavior of a random member of the previous generation. After each generation is born, the environment changes state with chance *u*, rendering all previously learned behavior non-adaptive. Therefore whether or not social learners acquire adaptive behavior depends upon both the strategy of the individual they copy as well as how long ago the environment changed state. This much is merely Rogers’ model with the addition of a chance 1 — *s* that innovation fails.

Now to these assumptions, add another dimension of behavior. In addition to being *adaptive* or *non-adaptive,* learned behavior can be simultaneously either *parasitic* or *non-parasitic.* This implies four types of behavior that may coexist in the population: adaptive/parasitic, adaptive/non-parasitic, non-adaptive/parasitic, and non-adaptive/non-parasitic. Each of these combinations entails a specific expected marginal fitness for its host:

|  | adaptive | non-adaptive |
| --- | --- | --- |
| parasitic | $b - v$ | $-v$ |
| non-parasitic | $b$ | $0$ |

Any parasitic behavior, regardless of whether or not it is also adaptive, reduces its bearer’s fitness by *v*.

But parasitic behavior also increases the odds, by a factor *w >*1, that its bearer is copied by a social learner. Specifically, let *r* be the frequency of parasitic behavior in the parent generation. Let *R* be the probability a social learner copies an adult with parasitic behavior. Then the odds *R*/(1 — *R*) that a social learner in the offspring generation copies an adult with parasitic behavior are given by:

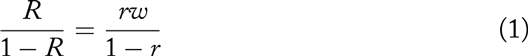

Solving for *R*:

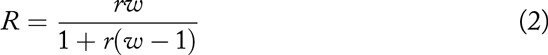

This reduces to *r* in the event that *w*=1, indicating unbiased social transmission. But when *w*>1, parasitized hosts are more influential.

This formulation has the advantage of being very general. It is a statistical description of a transmission advantage that may arise through diverse mechanisms. It has the disadvantage of implicitly assuming that the population is unstructured, so that a rare parasitized adult can potentially influence any individual in the population. As long as *w* is not too large, this assumption is perhaps not too odd, because the influence of a single individual will be very small. But for very large transmission advantage, a properly structured population model would be more useful.

### 2.2. Model dynamics

The assumptions above are sufficient to define the joint population dynamics for all four combinations of adaptive and parasitic behavior, as well as for the alleles for individual (I) and social (S) learning. To understand the implications of introducing parasitic behavior, I solve the dynamical system for its different steady states and the conditions that lead to each. The protect the innocent, a detailed derivation is presented only in the appendix.

Parasitic behavior invades and persists in this model when the frequency of social learning exceeds 1/*w*. When this holds, parasitic variants spread faster than individual learning introduces newly adaptive and non-parasitic variants. Why? Social learning influences the amount of parasitic behavior in the population, in two ways. First, parasitic behavior can only spread to social learners. Second, innovation tends 150 to be disassociated with parasitic behavior. When social learning is rare, innovation is common, and more non-parasitic behavior flows into the population each generation. Therefore when social learning is common, the rate of innovation is low and there are many potential hosts for parasitic behavior to spread to. As long as the transmission advantage *w* is large enough, and social learning is common enough, parasitic behavior can increase from zero and persist.

While parasitic behavior can easily invade, it has trouble becoming prevalent. Its prevalence is ultimately constrained by natural selection on social learning. In this model, the long-run fitness of social learning is constrained to equal the average fitness of individual learning. Why? Because social learning has highest fitness when it is rare. As it increases in frequency, its fitness declines. This continues until a long-run steady state is reached at which the expected fitness of social learning equals the expected fitness of individual learning. As parasitic behavior invades and reduces the fitness of social learners, this will in turn reduce the frequency of social learners until their average fitness is again equal to that of individual learning. This reduction in social learning in turn reduces the amount of parasitic behavior. Parasitic behavior can invade and persist, but it cannot effectively exploit the population.

This feedback is complex, so it is perhaps best captured by a visualization of the model’s dynamics. Figure 1 displays a single simulation, using discrete environmental changes, which result in sudden changes in the state variables. The horizontal axis is time, measured in generations. The top plot shows the correlation between adaptive and parasitic behavior in each generation. The bottom plot shows the frequencies of social learning (black), adaptive behavior (blue), and parasitic behavior (red) in each generation. At the start of the simulation, the frequencies of social learning and adaptive behavior are set to their steady state expectations in the absence of parasitic behavior. The proportion of social learning is high, just over 0.9, and the proportion of adaptive behavior is 0.3. Parasitic behavior is introduced into the population at a rate of 1 in 1000 innovation attempts. After several generations, it rapidly increases in frequency until the cost *v* begins to reduce the fitness of social learning. This leads to a decline in the frequency of social learning and an immediate reduction of the prevalence of parasitic behavior. As the frequency of social learning declines, the proportion of adaptive behavior in the population rises until the fitness of social learning again equals that of individual learning. At this point, parasitic behavior is maintained, but at a lower frequency than it initially achieved.

**Figure I.**
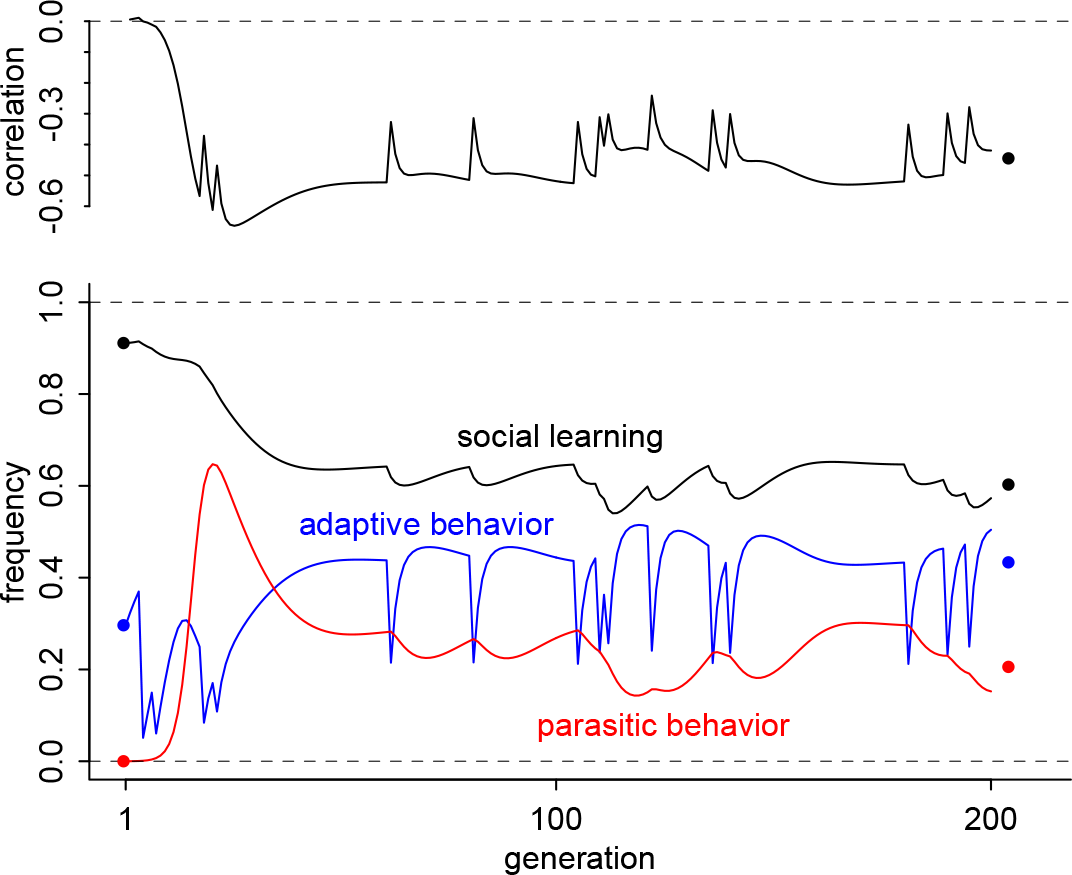
Dynamics of parasitic behavior, invasion and steady state. This plot displays a single numerical example of the model in this section, with parameters set so that parasitic behavior (shown in red) can invade the population. Top: Correlation between parasitic behavior and adaptive behavior. Bottom: Frequencies of social learning (black), adaptive behavior (blue), and parasitic behavior (red). The solid points in the right margin show the analytic expected values at steady state. After the initial outbreak, natural selection reduces social learning, which in turn reduces the prevalence of parasitic behavior. Ironically, the expected amount of adaptive behavior is increased, because there is more innovation. Parameter values for this example: *b/c* = 3, *s* = 0.6, *u* = 0.1, *v* = 0.5, *w* = 2, *w*_0_ = 10, *μ* = 0.001. Simulation code for this example is available at github.com/rmcelreath/parasiticbehaviorsim. Expressions for the analytical approximations (dots in the right margin) are found in the appendix.

Ironically, the proportion of adaptive behavior in the population is now higher than it was before parasitic behavior invaded. This is because the amount of innovation is now higher. But the population growth rate has not increased as a result,because the growth rate at steady state is still constrained to equal that of individual learning in isolation.

This model lacks explicit dynamics for the virulence, *v*, of parasitic behavior. However, since there is no population structure, as long as *w* is an increasing function of *v*, competition among parasitic variants will increase *v*. This will in turn reduce the frequency of social learning in the population, leading both to the collapse of parasitic behavior—a tragedy of the parasite commons—as well as an end to social learning, at least until the parasitic variants are eliminated and the process can begin again. I present a mathematical version of this argument in the appendix.
This is a logical scenario, but behavioral and psychological constraints on *v* and *w* may well limit its relevance. It is instructive only as it illustrates the fundamental conflict between parasitic behavior and social learning in this model.

## 3. STRATEGIC SOCIAL LEARNING AND PARASITIC BEHAVIOR

The model explored above builds on a venerable model structure. This makes it useful, because it is comparable to many other existing models. It sheds light on the fundamental question of what happens when parasitic behavior is introduced into a simple socially learning population.

But social learning, whether of humans or other animals, is frequently *strategic*, employing various cues to strategically select and transform socially learned behavior (Boyd and Richerson 1985, Henrich and McElreath 2003, Laland 2004). One consequence is that strategic social learning can produce a fitness surplus, increasing the frequency of adaptive behavior in the population above the raw asocial innovation rate—the *s* parameter in the previous model. Since much of the behavior in the previous model rested on the fact that, at steady state, social and individual learning had the same expected fitness, the dynamics of parasitic behavior in the context on strategic social learning may be rather different.

In this section, I construct and analyze a model comprising one mechanism of strategic social learning. I present another mechanism in the appendix.

### 3.1. Critical social learning

One way to make social and individual learning work synergistically is for the organism to deploy each when it is most useful (Boyd and Richerson 1995). Boyd and Richerson (1996) studied a strategy that used individual learning conditionally, only after failing to find a useful behavior by social learning. Enquist, Eriksson, and Ghirlanda (2007) analyzed a very similar strategy in more depth, under the name *critical social learning.* I’ve previously analyzed an unpleasantly complicated model of contingency to environmental change that ended up showing similar dynamics as critical social learning (McElreath 2012), suggesting that critical social learning serves a useful analytical role as one of the simplest and easiest to understand examples of conditional social learning. All of these analyses suggest that critical social learning, whatever we call it, can exclude both pure individual and pure social learning, raise the prevalence of adaptive behavior beyond the asocial innovation rate, and increase mean absolute fitness. It is also easy to do algebra with. This makes it a more interesting partner for parasitic behavior.

Define *critical social learning* as a heritable learning strategy that acquires candidate behavior by social learning and then evaluates the learned behavior, at a fitness cost h. When the candidate behavior is currently adaptive, the individual recognizes this fact with probability 1 — e. If instead the candidate behavior is currently non-adaptive, the individual detects this with probability *d*. If the individual rejects the candidate behavior, it attempts to innovate a currently adaptive behavior. This innovation entails a fitness cost *c* and succeeds with probability *s*, just as in the previous model.

These assumptions are sufficient to define a fitness expression for critical social learning and recursions for the population dynamics of all four combinations of adaptive and parasitic behavior. These expressions can then be solved for steady states and their stability conditions. Again to protect the innocent, I present the mathematical details only in the appendix.

This model produces a range of dynamics, from excluding parasitic behavior to extensive epidemics to mutualistic interactions between social learning and parasitic behavior. I summarize these dynamics by explaining four dynamic regimes:

1. *No parasites:* Just like in the previous model, it is possible for the population to exclude parasitic behavior.
2. *Persistent epidemic:* When parasitic behavior invades, it may become associated with non-adaptive behavior, reduce the level of adaptive behavior, and reduce host fitness. This epidemic persists across changes in the environment, although it is hard for it to spread to the entire population.
3. *Extensive epidemic:* Parasitic behavior may invade, but fail to reduce the level of adaptive behavior. This allows it to spread rapidly and to most of the population. It may substantially reduce host fitness.
4. *Mutualistic epidemic:* Parasitic behavior may invade, become positively associated with adaptive behavior, raise the level of adaptation, and substantially increase host fitness.

### 3.2. No parasites

I show in the appendix that the condition for critical social learning to exclude parasitic behavior is that both (1 — *d*) and (1 — *u*)(1 — *e*) must be less than 1/w. The intuition behind this condition is that there are two paths for parasitic behavior to spread within the population: either linked to adaptive or rather to non-adaptive behavior. To prevent the spread of parasitic behavior, both paths must be blocked. Parasitic behavior has an advantage *w* in becoming a candidate behavior. Then it is evaluated, and depending upon whether the behavior is also adaptive, different processes come into play. When linked to non-adaptive behavior, candidate parasitic behavior will be rejected *d* of the time, when the individual correctly detects non-adaptive behavior. When instead parasitic behavior is linked to adaptive behavior, it may spread and persist only if it is not rejected as non-adaptive, which happens *e* of the time. If it passes that filter, then the environment must not change, as happens 1 — *u* of the time. When the environment changes, all parasitic behavior is immediately linked to non-adaptive behavior, sending it into the other pathway in which *d* is relevant.

**Figure 2.**
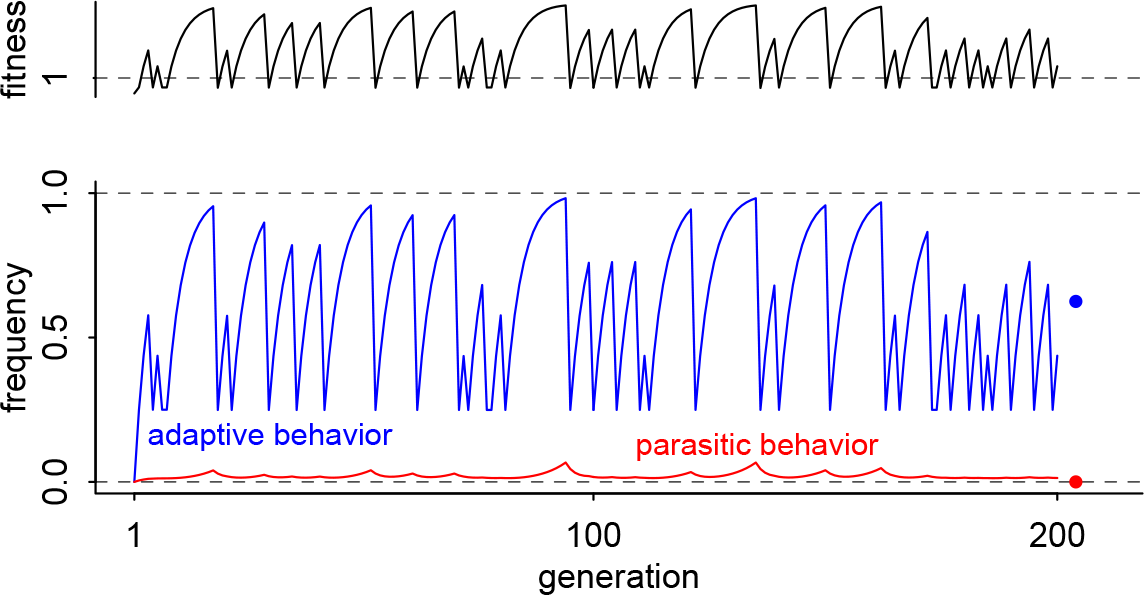
Dynamics of the critical social learning model, when parasitic behavior cannot invade. Top: mean fitness relative to pure individual learning. Bottom: frequencies of adaptive behavior (blue) and parasitic behavior (red). Analytical expected values are shown by the points in the righthand margin. Even with a very high innovation rate (*μ* = 0.01), parasitic behavior cannot invade, provided *w*(1 — *u*)(1 — *e*) and *w*(1 — *d*) are both less than 1. Parameter values for this example: *b/c* =3, *s*= 0.5, *u* = 0.2, *d* = 0.5, *e* = 0, *h* = 1/20, *v* = 0.5, *w* = 1.2, *w* = 10, *n* = 0.01. Simulation code for this example is available at github.com/rmcelreath/parasiticbehaviorsim. Expressions for the analytical approximations (dots in the right margin) are found in the appendix.

It will be useful to take a look at the behavioral dynamics of this model, when parasitic behavior is excluded. Unlike the previous model, the frequency of adaptive behavior can climb above the innovation rate *s*. This is because detection of non-adaptive behavior and retention of adaptive behavior ratchets the frequency of adaptive behavior up across generations and between changes in the environment. Figure 2 presents a representative simulation. In this example, a strong innovation rate of parasitic behavior (*μ* = 0.01) constantly introduces parasitic variants (red). But because critical learning is efficient and the environment changes frequently, parasitic behavior cannot persist. Adaptive behavior (blue) however is able to ratchet up to very high frequencies, despite the innovation rate of adaptive behavior being only *s* = 0.5. Because the frequency of adaptive behavior regularly exceeds s, the population enjoys a mean fitness in excess of pure individual learning, as shown in the top panel.

### 3.3. Persistent epidemic

When parasitic behavior can invade and persist, several different regimes may arise, depending upon the balance of forces. The first regime to consider is a persistent epidemic that resembles the epidemics from the simple social learning model. This regime arises when critical learning is not very accurate, relative to the rate of environmental change.

In this regime, a reliably negative covariance develops between parasitic and adaptive behavior. In fact, the analytical expression for the expected covariance (presented in the appendix) is identical to that of the previous model. The reason the covariance is reliably negative is partly the same as in the previous model: when the environment changes, all previously adaptive behavior becomes non-adaptive. All else being equal, this tends to associate parasitic behavior with non-adaptive behavior over time. However, in this model there is also a filtering process: critical social learners more often reject non-adaptive than adaptive behavior. Since parasitic behavior has a transmission advantage, it has greater chances of being evaluated. Since non-adaptive/parasitic combinations are rejected more often than adaptive/parasitic combinations, the critical learning filter can, given enough time, associate parasitic behavior with adaptive behavior. Is there enough time? When the environment changes quickly, relative to the detection probability *d*, there is not enough time.

The regime that results, when the rate of change *u* overpowers critical learning *d*, produces epidemic parasitic behavior that may severely depress mean fitness of the host. But the epidemic also hurts the behavioral ratchet of critical social learning, and this too limits the extent of the epidemic. Parasitic behavior is persistent, but it often fails to spread to the entire population. Figure 3 demonstrates this regime for *d* = 0.3 and *u* = 0.2 (other parameter values in the figure caption). The population enjoys a brief period of widespread adaptive behavior (blue), before parasitic behavior (red) invades. Afterwards, the frequency of adaptive behavior is suppressed by the negative correlation between parasitic and adaptive behavior—social learning is guided towards too much non-adaptive behavior, and critical learning is not accurate enough to counteract the transmission bias.

However parasitic behavior is also limited under this regime. Instead of continuing to spread throughout the population, it reaches a plateau frequency and remains there, as seen in the example in Figure 3. If *w* is very large, this plateau can be very high, but it still exists. Why doesn’t parasite behavior continue spreading throughout the entire population? When parasitic behavior is associated with non-adaptive behavior, it spreads mainly through the non-adaptive pathway, through not being rejected by critical social learning, 1 — *d* of the time. This means there is a constant and sizable force reducing parasitic behavior in the population.

### 3.4. Extensive epidemic

When *d* is instead sufficiently greater than *u*, the population dynamics change dramatically. Because *d* is relatively large, the best way for parasitic behavior to spread is by attaching itself to adaptive behavior. The first order effect of this is that parasitic behavior does not limit the spread of adaptive behavior. Instead the population of social learners can achieve very high frequencies of adaptive behavior. The second order effect is that parasitic behavior itself can spread just as high. If parasitic behavior is sufficiently costly, this can erase any fitness gains from higher levels of adaptive behavior. Figure 4 displays an example of this regime. Notice, in the top two panels of Figure 4, that the correlation between adaptive and parasitic behavior (middle panel) reliably evolves upwards, between changes in the environment. This positive covariation is what allows parasitic behavior to spread, because it effectively bypasses the skepticism of critical social learners. As parasitic behavior rises in frequency, mean fitness (top panel) declines, stopping just above 335 that of pure individual learning, shown by the dashed line.

**Figure 3.**
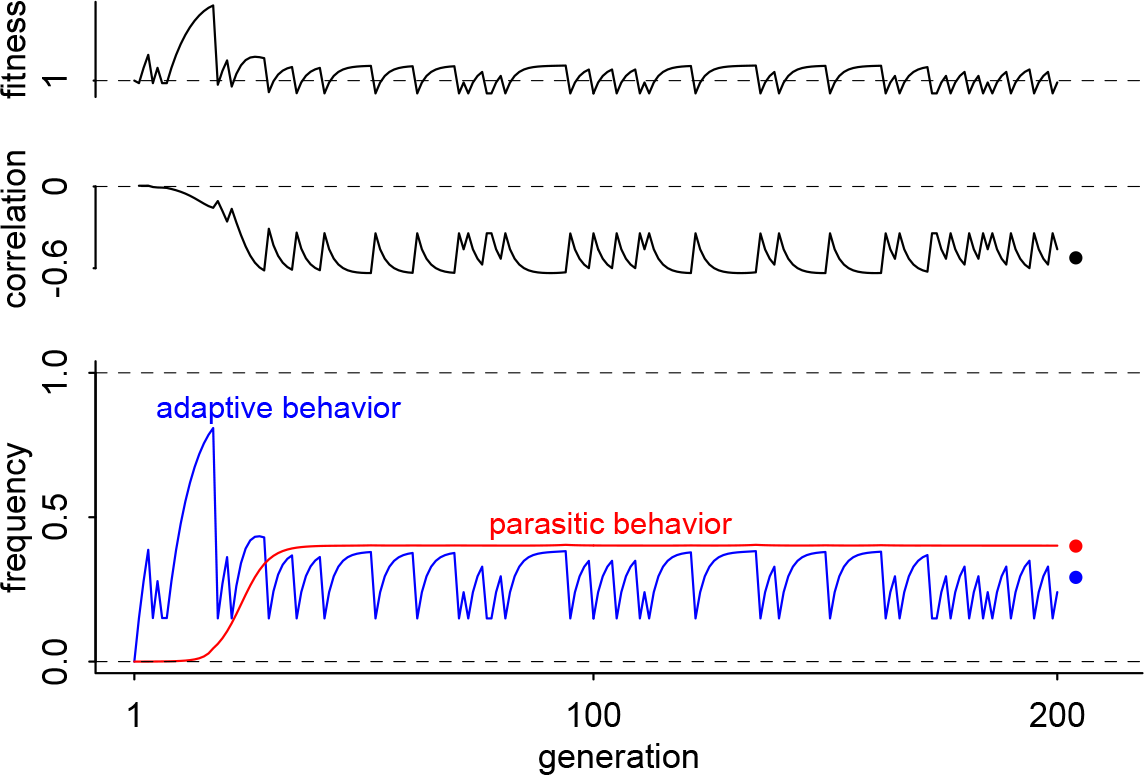
When parasitic behavior becomes associated with non-adaptive behavior. From top to bottom: mean fitness relative to pure individual learning, correlation between adaptive and parasitic behavior, frequencies of adaptive (blue) and parasitic behavior (red). Before parasitic behavior invades, adaptive behavior can ratchet above the innovation rate *s*. Once it invades, parasitic behavior suppresses adaptive behavior. Parameter values for this example: *b/c* = 2, *s* = 0.5, *u* = 0.2, *d* = 0.3, *e* = 0, *h* = 1/20, *v* = 0.5, *w* = 2, *w*_0_ = 10, *μ* = 10^−4^. Simulation code for this example is available at github.com/rmcelreath/parasiticbehaviorsim. Expressions for the analytical approximations (dots in the right margin) are found in the appendix.

While the example in Figure 4 does not result in the loss of social learning, this is possible with sufficiently high *v*. By making *w* and *v* sufficiently large, the population will suffer the partial loss of social learning, as pure individual learners who are immune to parasitic behavior invade. This leads to the loss of parasitic behavior, 340 which is then followed by the re-invasion of critical social learning, and the cycle begins anew. I encourage readers to download the simulation code and conduct their own experiments.

**Figure 4.**
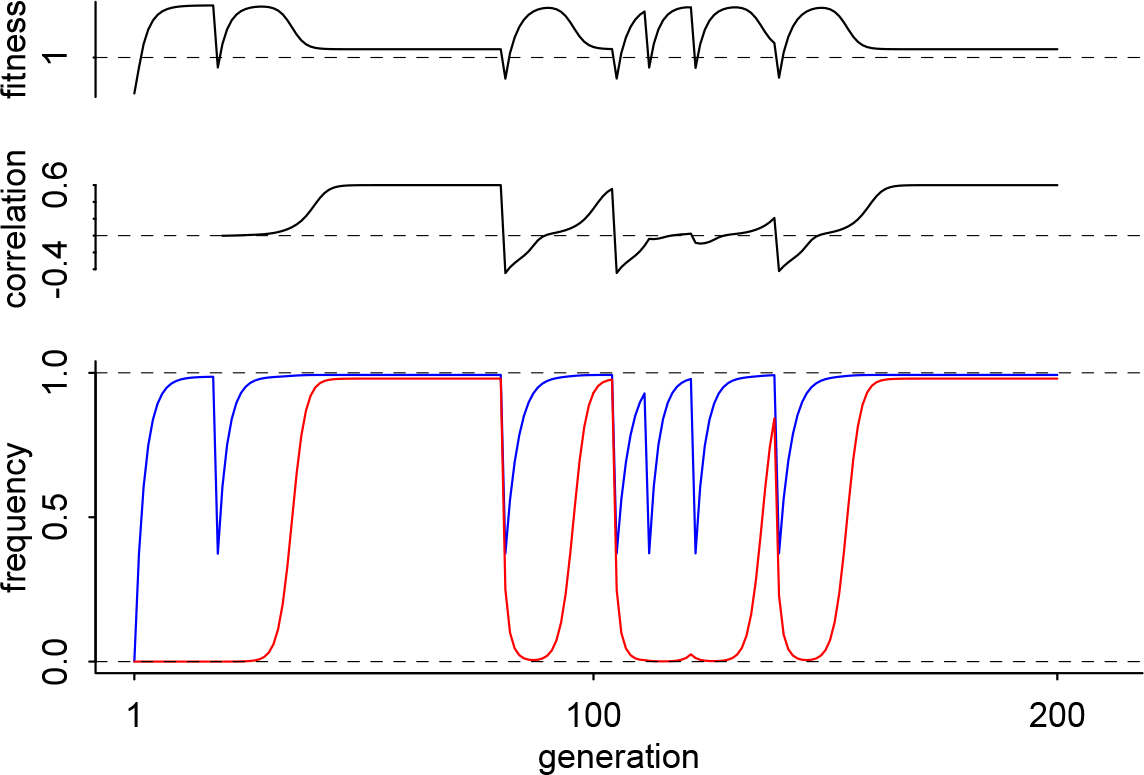
When parasitic behavior invades and results in an extensive epidemic. From top to bottom: mean fitness relative to pure individual learning, correlation between adaptive and parasitic behavior, frequencies of adaptive (blue) and parasitic behavior (red). Before parasitic behavior invades, adaptive behavior can ratchet above the innovation rate s. But once it invades, parasitic behavior suppresses adaptive behavior, due to its persistently negative covariance. Parameter values for this example: *b/c* = 10, *s* = 0.5, *u* = 0.05, *d* = 0.75, *e* = 0.01, *h* = 1/10, *v* = 5, *w* = 2, *w*_0_ = 10, *μ* = 10^−4^. Simulation code for this example is available at github.com/rmcelreath/parasiticbehaviorsim.

### Mutualistic epidemic

But when the cost imposed by parasitic behavior, *v*, is not too large, and the ceiling for adaptive behavior in the absence of parasitic behavior is sufficiently low, the spread of parasitic behavior can actually propel the further spread of adaptive behavior and increase the mean fitness of the host. This regime is demonstrated in Figure 5. In this example, *d* is again large relative to *u*. As a result, parasitic behavior must again spread predominantly through being linked to adaptive behavior. But now critical social learning is also prone to false positives, *e* = 0.1. This imposes a ceiling on how far adaptive behavior can spread in the population, as seen in the dynamics before parasitic behavior invades. Adaptive behavior levels off before reaching a frequency of 0.5.

After parasitic behavior invades, however, a positive covariance (middle panel, Figure 5) again evolves. The transmission advantage of parasitic behavior now helps to spread adaptive behavior, counteracting the downward force of false positive errors e. Both adaptive and parasitic behavior can reach high frequencies, before the environment changes state and the process begins again. The result is that mean fitness (top panel) can be substantially increased, as long as the cost of parasitic *v* is not too large.

**Figure 5.**
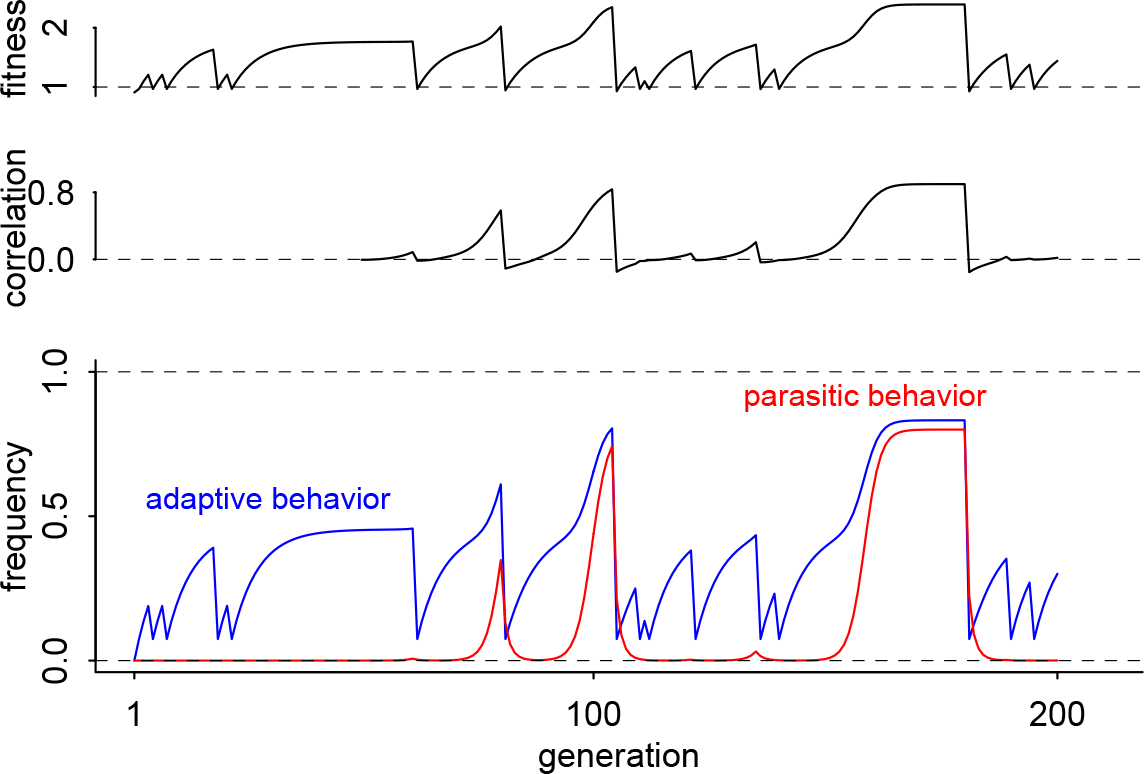
When parasitic behavior invades and increases host fitness. From top to bottom: mean fitness relative to pure individual learning, correlation between adaptive and parasitic behavior, frequencies of adaptive (blue) and parasitic behavior (red). Before parasitic behavior invades, learning errors (e, 1 — *d)* and a low innovation rate *s* limit adaptive behavior. After parasitic behavior invades, it evolves a positive correlation with and guides individuals to adaptive behavior. The population does best when parasitic behavior is common. Parameter values for this example: *b/c* = 20, *s* = 0.1, *u* = 0.1, *d* = 0.75, *e* = 0.1, *h* = 1/10, *v* = 2, *w* = 2, *w*_0_ = 10, *μ* = 10^−4^. Simulation code for this example is available at github.com/rmcelreath/parasiticbehaviorsim.

## 4. LEARNING, BEHAVIOR, AND DANGEROUS THINGS

The goal of these analyses is nothing more than to begin to explore the coevolution of social learning and the potential for parasitic behavior. To that end, I have developed and analyzed two models. These models are very general, taking structurally familiar models of the evolution of social learning and introducing parasitic behavior. The models do not assume social learning and ask whether parasitic behavior can spread. Instead they simultaneously allow for the possibility of social learning and parasitic behavior and derive when either or both can spread.

### 4.1. Summary of the First Model

The first model adds one kind of parasitic behavior to the most common and simplest structure for exploring the evolution of social learning (Rogers 1988). All that has been added is the possibility of behavior that imposes a fitness cost on its carrier and thereby increases its odds of being transmitted to the next generation of social learners. Such behavior may also be adaptive. This model is among the most minimal models of the coevolution of social learning and parasitic behavior that can be devised. As such, it is both deeply unsatisfying and a necessary step towards more sophisticated arguments.

In an evolutionary narrative, this model stands in for a population in which primitive social learning has emerged and must confront the risk of spreading parasitic behavior. My analysis suggests that the courtship between primitive social learning and parasitic behavior is ultimately frustrating for both: social learning is counter-selected in the presence of parasitic behavior and parasitic behavior can never be come very common in such a population.

Really this coevolutionary response arises from the unbiased nature of social learning. Social learning in this model can never produce a surplus for the population, because it does no more than mirror the frequency of adaptive behavior among individual learners. But the important implication is that any damage that parasitic behavior does to social learners will result in fewer social learners at steady state. Parasitic behavior cannot exploit social learners without also reducing their numbers.

### 4.2. Summary of the second model

The second model behaves quite differently. In models with strategic social learning strategies, social learning is not just a form of social loafing but may actually increase absolute mean fitness. In that case, parasitic behavior can do a lot more, or a lot less, damage.

To appreciate why it might do more damage, consider that the fitness surplus strategic social learning produces provides an evolutionary buffer. This buffer protects social learning against competition from individual learning, once parasitic behavior invades. The cost of practicing parasitic behavior must exceed the strategic surplus, before natural selection will reduce the amount of social learning and therefore limit the spread of parasitic behavior. In one sense, the worst that can happen is that the mean fitness is again limited to that of pure individual learning, as in the simple model. But compared to what could be achieved in the absence of parasitic behavior, large amounts of fitness—and therefore potential population growth—can be lost.

Beyond considerations about population growth, strategic social learning allows parasitic behavior to reach much higher prevalence. This drastically changes predictions about the adaptive nature of socially transmitted behavior. Ironically, it is when social learning is more sophisticated that common behavior may sometimes be, as is often suggested, not for the organism but rather for itself.

However strategic social learning does not necessarily result in such a morose outcome. I showed that critical social learning (Boyd and Richerson 1996, Enquist et al. 2007) may also lead to mutualistic interaction between social learning and parasitic behavior. Parasitic behavior becomes positively associated with adaptive behavior, which allows it to spread through a population of selective learners. Social learners in turn benefit from the fact that parasitic behavior then advertises adaptive behavior. Despite the fitness cost this advertisement imposes on social learners, the interaction brings a net fitness benefit. It also brings a benefit to parasitic behavior, as it is able to achieve a much higher prevalence in the presence of critical social learning.

Would other forms of strategic social learning lead to similar dynamics? I have also analyzed a model of success-biased social learning in the presence of parasitic behavior. I present this model very briefly in the appendix. This model may also produce a positive covariance between adaptive and parasitic behavior. And under some circumstances, this positive covariance can increase mean fitness. It may also result in terrible epidemics.

Similar dynamics arise in the success-bias case for two reasons. First, strategic social learning again allows the population to sustain much more social learning than would be possible under simple, unbiased modes of transmission. This allows parasitic behavior to become prevalent. Second, the inferential force of success-biased learning, like critical social learning, is to favor combinations of parasitic and adaptive behavior. This makes a positive covariance possible and even likely under the right circumstances. Because diverse forms of strategic social learning should support both of these phenomena, the general result many persist under many different mechanisms.

I caution however that the exact dynamics of the success-bias model are rather different from those of the critical social learning model. Therefore they are likely to interact with other learning strategies and population structures in rather different ways. Other forms of strategic social learning, such as conformist transmission (Boyd and Richerson 1985), could be even more different, especially as conformist transmission interacts quite differently with spatial versus temporal environmental variation (McElreath et al. 2013, Nakahashi et al. 2012).

### 4.3. The natural selection of psychological vulnerability

In the course of this work, I have assumed that parasitic behavior is unavoidable. This follows from the motivating hypothesis: Social learning brings with it some risk of manipulation. Just like breathing entails exposure to pathogens, social learning entails exposure to manipulation, either by other individuals or by behavior itself.

But when “manipulative” behavior successfully guides a learner to adaptive behavior, the situation seems less manipulative and more mutualistic. Thus under the right circumstances, it may be evolutionarily advantageous for a social learner to be susceptible to parasitic behavior. In the critical social learning model, an alternative learning allele that were less susceptible to parasitic variants—experiencing a lower effective probability of learning a parasitic variant—would have have lower fitness and be unable to invade a population of ordinary, susceptible critical social learners, provided the covariance between adaptive and parasitic behavior were high enough to compensate for the fitness cost of parasitic behavior. I present an analytical condition in the appendix.

This insight may not be of general value. But it offers another way to understand cultural cognition. Instead of parasitic behavior arising merely from the inherent tradeoff that social learning requires credulity, it makes that very credulity appear useful to the organism. It reflects a kind of evolutionary rationality that makes sense only in the context of particular behavioral environments constructed and maintained by the mass action of seemingly irrational individuals (McElreath et al. 2013).

### 4.4. Limitations and extensions

Many good theories lack nuance (Healey 2016). The models I have contributed in this paper certainly lack nuance. This does not necessarily mean that they in turn contribute to good theory. But their lack of nuance should not be held against them. Nevertheless, there are several features of these models that deserve critical attention, without falling into any nuance traps. In closing, I discuss some prominent issues for follow-up investigation.

Population structure is very important in both social learning and disease transmission. The models herein have no population structure, being neither divided into finite groups nor having varying connectivity among individuals. Why might population structure matter? Consider that variation in connectivity has the main effect of reducing disease spread, while making rare outbreaks more serious (Lloyd-Smith et al. 2005). By analogy, the spread of parasitic behavior will depend strongly on network structure. Age structure is no less important, especially when combined with age-dependent migration.

Population structure is important for the virulence of behavior, as well. In these models, due to the absence of structure, parasites may evolve a tragedy of the commons in which they become too virulent for their own good. Population structure may alter this result, by creating the analog of kin selection, or equivalently group selection, among parasitic variants. Population structure will also change the nature of any transmission advantage, as no longer will the entire population of learners be influenced by the same pool of social models.

The models here assume oblique social learning between generations. Other patterns of social learning may induce different dynamics. Vertical transmission— learning from parents—may reduce the spread of parasitic behavior, but may also reduce incentives for strategic learning. Vertical transmission also interacts with effects on fertility and survival in different ways (McElreath and Strimling 2008), meaning that parasitic behavior may develop conditional correlations with particular fitness effects. True horizontal transmission within age classes might accelerate the spread of parasitic behavior, but it may also increase incentives for strategic learning in compensation.

I have neglected discussion and analysis of the innovation rate of parasitic behavior, usually assuming it is vanishingly small. This is a strategic assumption that highlights the forces central to the motivating hypothesis. But innovation and transmission do interact to determine dynamics. In the critical social learning model, increasing the innovation rate of parasitic behavior actually makes it easier to achieve a positive covariance between adaptive and parasitic behavior. Why? Because when parasite innovations are rare, this makes the initial covariance between adaptive and parasitic behavior more negative. As more initial combinations of adaptive and parasitic behavior are innovated, the covariance can build faster due to the transmission dynamics. This is especially important when the environment changes rapidly, and so there is limited time for the positive covariance to build. The reader can easily appreciate this innovation-transmission interaction by replicating the simulation in Figure 5 and gradually increasing the innovation rate *μ* (mu in the code). Getting analytical results for this interaction is difficult. Even using aggressive approximations, I have not succeeded. But I am confident that others might succeed or rather make effective study with numerical methods.

None of the models in this paper are the final word on anything. Including forces that have been omitted from these models would modify the conclusions. But the fact that even simple models with only simple selective forces can produce diverse and surprising results should make us hesitate and reevaluate previous verbal arguments. These models comprise an invitation to an ongoing discussion of the role of selection in explaining both the design of behavior and the nature of animal psychology.

## APPENDIX

**Recursions, fitness expressions, and steady state solutions for the simple social learning model**. Let *x*_11_ indicate the frequency of adaptive/parasitic behavior in the population, *x*_00_ the frequency of non-adaptive/neutral behavior, and *x*_10_ and *x*_01_ the frequencies of the mixed types. In addition to *p*, the frequency of social learning alleles in the population, these variables define the state of the population in each generation. Let *Q*_11_ be the probability a social learner acquires adaptive/parasitic behavior. Let *μ* be the probability that an act of innovation produces a new parasitic variant. Then the recursions for each *x_ij_* are:

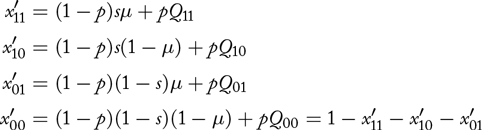

And the expressions for each *Qij* are:

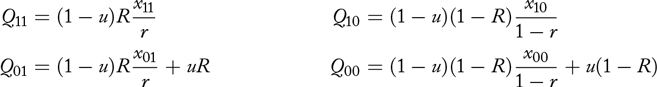

where *R* = *rw/*(*rw* + 1 — *r*) is the probability a social learner copies a parasitized individual, as derived in the main text. The expression for *Q*_11_ can be motivated by noting that it is only possible to acquire adaptive/parasitic (11) behavior when the environment has not recently changed. This happens 1 —*u* of the time. Then *R* is the probability a social learner selects a parasitized adult. Finally, *xn/r* is the conditional probability of acquiring adaptive behavior, conditional on the social model having parasitic behavior as well. The expression for *Q*_10_ is motivated similarly. The expressions for *Q*_01_ and *Q*_00_ contain additional terms starting with *u* to account for the conversion of adaptive/parasitic to non-adaptive/parasitic and adaptive/non-parasitic to non-adaptive/non-parasitic, when the environment changes.

It is more useful to analyze the system in terms of *q* = *x*_11_ + _10_, the frequency of adaptive behavior, *r* = *x*_11_ + *x*_01_, the frequency of parasitic behavior, and *k* = *x*_11_*x*_00_ — *x*_10_*x*_01_, their covariance. Substituting for each *x_ij_* in terms of *q*, *r*, and *k* yields these recursions, where *μ* → 0 for simplicity:

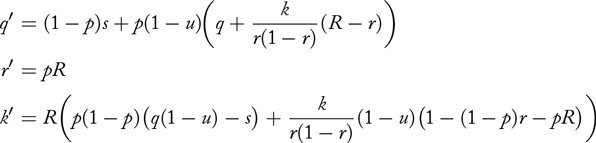

The recursion for*p* the frequency ofsociallearning (S), is given by the usual discrete-generation replicator equation:

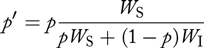

where *W_S_* = *W*_0_ + *b*(*Q*_11_ + *Q*_10_) — *v*(*Q*_11_ + *Q*_01_) is the average fitness of social learners and *W*_I_ = *w*_0_ + *sb* — *c*.

This is a stochastic system, and so it has no actual equilibrium. Instead, environmental fluctuations periodically change the selection gradient. Evolution in these models is therefore restless and even exciting. And like in classical host-parasite models, this model can produce oscillations and highly stochastic outbreaks and recoveries. Solutions are available however for the expected values of the state variables, assuming that selection is weak enough that the cultural dynamics are essentially at steady state for any given allele frequencies. This is the same separation of time scales approximation that is used in solving many models in this literature. The steady state solutions for the expected values of the state variables *p*, *q*, *r*, and *k* are found by taking the expectation of each recursion over time and then solving for values of the expected values of*p*, *q*, *r*, and *k* that remain unchanged from generation to generation. There are five sets of solutions. The solutions are given by:

| Steady state | $\hat{p}$ | $\hat{q}$ | $\hat{r}$ | $\hat{k}$ |
| --- | --- | --- | --- | --- |
| (1) Pure individual | 0 | $s$ | 0 | 0 |
| (2) Pure social | 1 | 0 | 0 | 0 |
| (3) Pure epidemic social | 1 | 0 | 1 | 0 |
| (4) Mixed | $\frac{1 - usb/c}{1 - u}$ | $\frac{1 - \hat{p}}{ub/c}$ | 0 | 0 |
| (5) Mixed epidemic | $\frac{sb/c(1 - u) - \frac{v}{c} \frac{(u+w-1)}{w-1}}{(u + w - 1)(1 - \frac{v}{c} \frac{w}{w-1}) - wsb/c}$ | $\frac{(1 - \hat{p})sw}{u + w - 1}$ | $\frac{\hat{p}w - 1}{w - 1}$ | $-\hat{q}\hat{r}$ |

The find when these solutions are each stable against perturbations in any dimension, I linearized the system of recursions in the neighborhood of each set of solutions and solved for the conditions that make the dominant eigenvalue less than one. The stability conditions for each of the five possible steady states are given by:

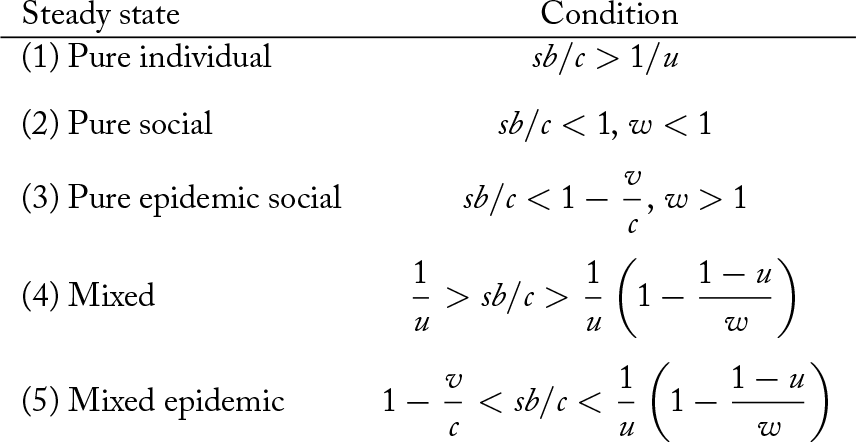

| Steady state | Condition |
| --- | --- |
| (1) Pure individual | $sb/c > 1/u$ |
| (2) Pure social | $sb/c < 1, w < 1$ |
| (3) Pure epidemic social | $sb/c < 1 - \frac{v}{c}, w > 1$ |
| (4) Mixed | $\frac{1}{u} > sb/c > \frac{1}{u} \left(1 - \frac{1-u}{w}\right)$ |
| (5) Mixed epidemic | $1 - \frac{v}{c} < sb/c < \frac{1}{u} \left(1 - \frac{1-u}{w}\right)$ |

Since steady state 2 is only stable when *w* < 1, and the model requires *w* > 1, it is not discussed in the main text.

The stability regions for four of these steady states that are possible when *w* > 1 (1 and 3—5) are shown in Figure 6. The four steady states each represent an outcome that can teach us about the coevolution of social learning and parasitic behavior. The parameter regions that stabilize each of these four steady states are shown by the colored regions in Figure 6. There are four parametric dimensions that define the dynamical boundaries in this model: *sb/c*, 1/*u*, *v/c*, and *w*. The figure shows the four steady states as regions within coordinates of*sb/c* and 1/*u*, for fixed values of *v/c* and *w*. These regions are bounded by mutually exclusive stability conditions.

The blue region in the upper left represents a steady state in which individual learning is evolutionarily stable. As a result, there is no social transmission and therefore no parasitic behavior. This steady state is stable when individual learning produces a larger average fitness return than rare social learning, when *sb* — *c* > *sb*(1 — *u*). As long as the expected benefit of individual learning, *sb/c*, exceeds the expected waiting time until a change in the environment, 1/*u*, social learning cannot invade. This is a classic result found in many models.

The second region, shown in green, represents a stable mix of individual and social learning. But this mix may resist the invasion of parasitic behavior, provided the frequency of social learners is not too high. This state of the population is stable when:

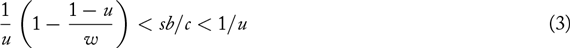

In plainer language, individual learning cannot be so profitable as to exclude social learning, but social learning cannot be so common that it spreads parasitic behavior faster than individual learners discover adaptive behavior. The relationship between the spread of parasitic behavior and the frequency of social learning can be deduced from the recursion for *r*, the proportion of behavior that is parasitic:

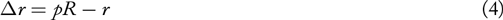

And so *r* cannot increase when *p* < *1/R* = (*rw* + 1 — *r)/w*. When parasitic behavior is rare, this condition is approximately *p* < 1*/w*. As a result, when w is large, and so parasitic behavior has a great advantage in transmission, the frequency of social learning must be very small or else parasitic behavior will invade. Social learning will be relatively rare at the mixed steady state when the expected time between changes in the environment, 1/*u*, is small. Together, these relationships explain the condition for a mixture of individual and social learning to resist invasion of parasitic behavior.

The red region is perhaps the most interesting. It indicates a mix of individual and social learning with epidemic parasitic behavior. This steady state persists when there are sufficient social learners for parasitic behavior to invade 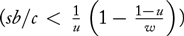 but individual learning is better than parasitized social learning 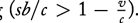. When both conditions hold, the population sustains both learning strategies as well as parasitic behavior. This region extends forever off the right side of Figure 6, as longer durations between environmental changes, 1/*u*, favor yet more social learning. However, unless an individual learner does worse than a parasitized social learner, individual learning will never be entirely removed from the population.

**Figure 6.**
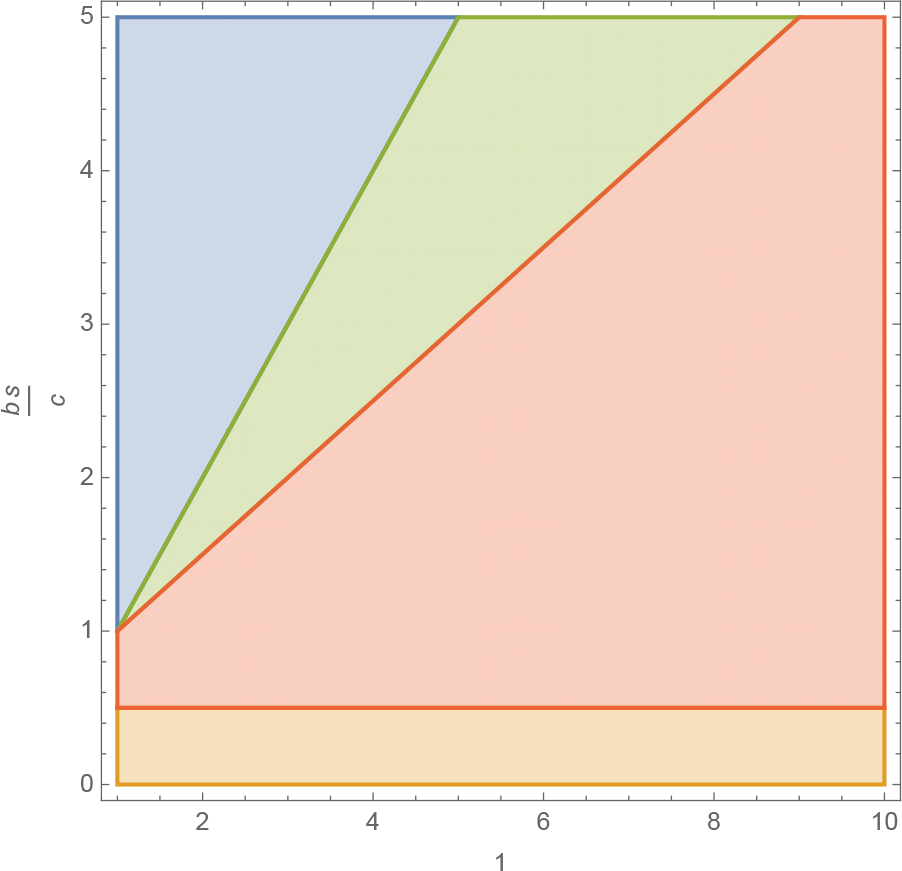
The four possible steady states ofthe simple parasitic behavior model. Each colored region defines the combinations of *sb/c* and 1/*u* that give rise to a unique steady state, for fixed values of *w* and *v*. Blue: Pure individual learning. Green: Un-parasitized mix of individual and social learning. Red: Parasitized mix of individual and social learning. Orange: Parasitized pure social learning.

Note that the condition for parasitic behavior to persist does not depend upon *v*, the fitness cost it imposes on an individual. However, *v* does indirectly influence the frequency of parasitic behavior, by reducing the fitness of social learning. The frequency of parasitic behavior at this steady state is given by:

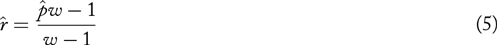

where 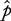 is the frequency of social learning. So the more selection favors social learning, the more extreme the behavioral epidemic. So when *v* is large, 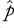 will be smaller at steady state, and the behavioral epidemic will be reduced.

The expected covariance between adaptive and parasitic behavior is given by:

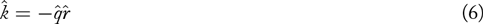

This is always negative, indicating that parasitic behavior ends up reliably associated with non-adaptive behavior. Why? Because when the environment changes state, all adaptive behavior is flushed. But parasitic behavior is not similarly flushed from the population. Then as individual learners innovate new behavior, only a small fraction of that will also be parasitic, relative to the persisting socially-transmitted parasitic behavior from before the change in the environment. Over time, parasitic behavior becomes associated with non-adaptive behavior. Part of the fitness cost it imposes on the host organism is through this dynamic mechanism, not just through the direct cost *v*.

The final region, at the bottom of the figure, corresponds to steady state in which individual learning is finally removed. The condition for this steady state is *sb/c* < 1 — *v/c*, which indicates that the relative fitness cost of parasitic behavior, *v/c*, is less than the relative cost of innovation. When this is true, social learning is evolutionarily stable and all behavior becomes non-adaptive and parasitic. This is perhaps a merely logical, but biologically irrelevant, steady state. It nevertheless helps us understand the model’s dynamics, as logical extremes often do.

**Adaptive dynamics of parasitic behavior virulence *v* in the simple social learning model**. Let the transmission advantage of parasitic behavior *w* be an increasing function of its virulence *v*, *w*(*v*\*). Imagine the entire population of parasitic variants have the same virulence *v*\* and therefor *w\** = *w*(*v*)*. Now also imagine a mutant parasitic variant with a different value *v* = *v*\* + δ and *w* = *w*(*v*), where δ is small enough so that δ^2^ ≈ 0. When does this mutant spread? This is equivalent to asking which value *v*\* cannot be invaded by *v*.

To define a fitness from the perspective a parasitic behavior, suppose that each social learning is influenced by a large number of social models *n*. Then the probability that the mutant is acquired by a social learner is:

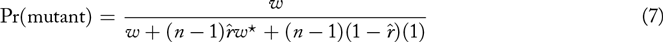

Essentially, the mutant gets *w* lottery tickets, while there are 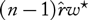 tickets from common-type parasites and (n — 1)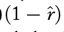 tickets from non-parasites. Note that the parasitic competition depends upon the prevalence 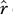 of parasitic behavior in the population, and this prevalence is determined by *w*\*, not by the mutant’s *w*.

There will be on average 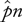 social learners in the population, and so 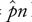 opportunities for the mutant to spread. This gives us the expected fitness of the mutant:

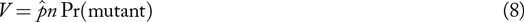

Taking the limit as *n*→∞:

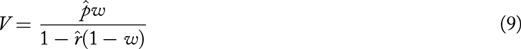

Now since 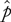 and 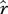 are functions of *w*\* and not *w*, it follows that *V* is strictly increasing with *w*. Therefore any mutant that satisfies *v* > *v*\* will invade.

It is also true that 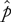 is a decreasing function of *v*, assuming *w*(*v*) is an increasing function of *v*. Therefore is also follows that as *v* increases in the population of parasitic variants, the steady state frequency of social learners will decrease. This is the tragedy of the commons, applied to parasitic behavior, where the “commons” means the social learning hosts. Group selected parasitic variants would presumably have a stable, intermediate value of *v* that maximizes prevalence.

**Recursions and fitness expressions for the critical social learning model**. Let *Q_ij_* represent the probability an individual acquires by social learning behavior *ij*, where *i* ∈ {0,1} indicates adaptive (1) or non-adaptive (0) behavior and *j* ∈ {0,1} indicates parasitic (1) or non-parasitic (0) behavior. Then the expected fitness of a critical social learner is:

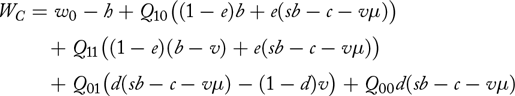

Note that the error rate *e*, relevant when adaptive behavior is acquired via social learning, and the error rate 1 — *d*, relevant instead when non-adaptive behavior is acquired, play analogous roles. A dissimilarity is that *e* can lead individuals to shed parasitic behavior, at a cost *c*, whereas the error from failing to discern non-adaptive behavior 1 — *d* of the time has no similar tradeoff.

The *Q_ij_* expressions are defined exactly as in the previous model, with a factor *w* > 1 that expresses the proportional odds of acquiring parasitic behavior. Let *pc* be the frequency of critical social learning, *pS* the frequency of pure social learning, and *pI* = 1 — *pC* — *pS* the frequency of pure individual learning. Let *Q* = *Q*_10_ + *Q*_11_ be the probability of acquiring adaptive behavior by social learning. Then the recursions for the proportions *x_ij_* of each combination of adaptive and parasitic behavior are however now:

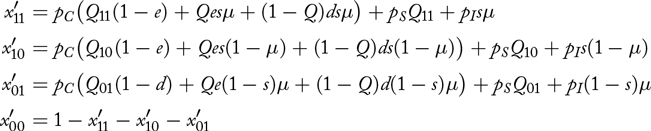

Again, it is more revealing to study the system in terms of the dynamics of *q* = *x*_11_ + *x*_10_, *r* = *x*_11_ + *x*_01_, and *k* = *x*_11_*x*_00_ — *x*_10_*x*_01_. These are complex expressions in this model, because they depend upon all three learning strategies and all four behavior combinations. But setting *pC* → 1 and *μ* →0, the recursions for *q* and *r* are (remember: *Q* = *Q*_11_ + *Q*_10_):

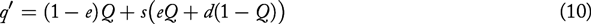

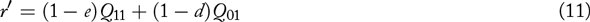

The *q* recursion reveals how the stock of adaptive behavior depends upon preserving adaptive behavior *Q* and the rate at which non-adaptive behavior is introduced. The recursion for *r* depends upon both adaptive and non-adaptive behavior. But adaptive behavior is potentially much more important, since 1 — *e* will usually be larger than 1 — *d*. This is to say the *r* depends upon how critical learning filters socially learned behavior. This dynamic ultimately explains why the covariance *k* can become positive and continue to grow.

To study the dynamics, assume weak selection so that we can separate the cultural and genetic time scales. Then we can solve for steady state expected values of *q*, *r*, and *k* in terms of *pC*, *pS*, and *pI*. These solutions can be quite complicated. In addition, they are much more approximate that in the previous model, because they ignore geometric fitness effects on the dynamics. The previous solutions also ignored such geometric effects, but now since the rate of individual learning is plastic, it changes on the cultural time scale and cannot really be fully separated from the cultural dynamics. The solutions to be derived this approximate way are still useful—simulation confirms that they provide useful approximations and valuable intuition. But they are not exact.

In expressions to follow, I set *pI* = 1 —*pC* — *pS*, *pC* = *p*, and *z* = *p* + *pS* for ease of notation. I also set *e* → 0, so that some of the expressions do not fill four lines. There are three sets of solutions, corresponding to three possible steady states:

1. No parasitic behavior

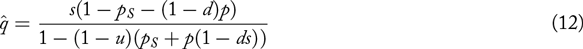

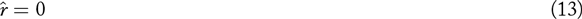

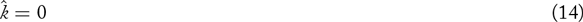
2. Epidemic Type I (Negative Covariance)

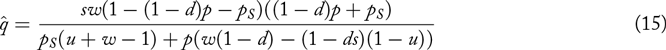

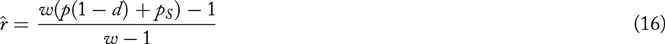

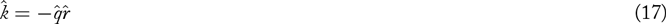
3. Epidemic Type II (General Covariance)

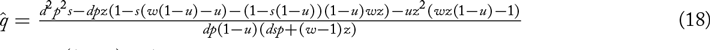

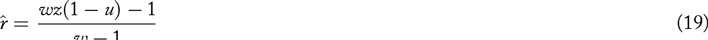

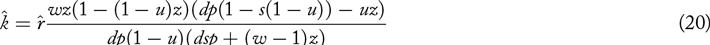

Note that the covariance 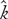 in the third solution will be positive whenever the term *dp*(1 − *s*(1 − *u*)) − *u* > 0. When critical social learning is an ESS, this condition becomes *d*(1 − *s*(1 − *u*)) − *u* > 0. Then as long as *d* is large enough, *s* is small enough, and *u* is small enough, the covariance between adaptive and parasitic behavior has a positive expected value.

The general conditions for each of these steady states to be stable are very complex. But when critical social learning dominates the population, the conditions are exceedingly simple. I derive them by linear stability analysis, yielding these sets of eigenvalues for each of the three solutions, respectively:

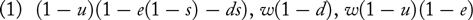

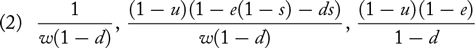

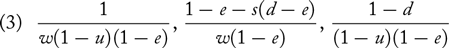

When all three expressions are less than 1, a solution is stable. Keep in mind that because this system is stochastic, these linear stability solutions are only approximations. They are useful and capture the economic tradeoffs in the model, but they do not provide exact boundaries for the behavior of the model. This is because there can be substantial geometric mean fitness effects as well as transient dynamics that are important.

To get some intuition for the transitions among these steady states, consider the first steady state, in which parasitic behavior is absent. To make the lesson exceedingly simple, also assume that critical social learning dom-inates the population (*pC* = 1) and that *e* = 0. When can parasitic behavior increase in frequency? The first parasitic variant can be paired with either adaptive behavior or non-adaptive behavior. Suppose it is paired with adaptive behavior. Then the change in *r* is approximately:

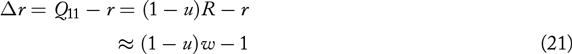

This is greater than zero when *w*(1 − *u*) >1. This is one of the sufficient conditions for parasitic behavior to invade. Now consider when the first parasitic variant is paired instead with non-adaptive behavior. Now the change in *r*is given by:

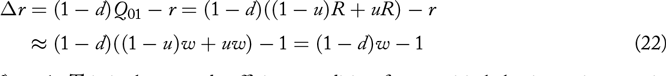

So *r* increases when *w*(1 − *d*) > 1. This is the second sufficient condition for parasitic behavior to increase in frequency. Both of these conditions can also be derived by linear stability analysis, of course, which is how I first found them. But the above derivation, connected directly to the recursion for *r*, is much more intuitive.

Once parasitic behavior invades, whether the system is attracted to the second or rather third steady state depends largely upon the relative sizes of *u* and *d* and *e*. Again assuming *e* = 0, the relevant comparison is
whether *d* > *u* or rather *u* > *d*. When *u* > *d*, then critical social learning is ineffective relative to the rate of environmental change, as explained in the main text. When instead *d* > *u*, the reverse is true. If *s* is also small enough, it is possible for critical social learning to effectively filter parasitic behavior and create a positive covariance between adaptive and parasitic behavior.

**Invasion by parasite-resistant social learners**. Under some circumstances, a mutant critical social learner who is resistant, or immune, to parasitic variants cannot invade the population. Suppose a completely immune mutant critical social learner C* for whom the probability of acquiring parasitic behavior is exactly its population frequency *r*. The fitness expression for this mutant is derived by substituting *R* → *r* in the fitness expression for ordinary critical social learners defined earlier. Subtracting the expected fitness of the invader from the expected fitness of the common type yields:

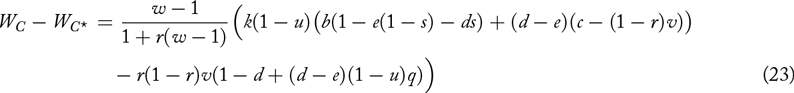

This is greater than zero, and so the mutant cannot invade, provided:

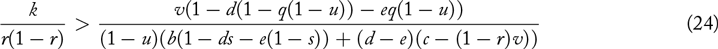

The left side is the regression slope predicting adaptive behavior, conditional on parasitic behavior. This is because a regression slope is the covariance between two variables divided by the variance in the conditioning variable. Therefore the left side states how well parasitic behavior predicts adaptive behavior. The right side is a terrifying calamity of expected marginal fitness effects. But it is clear that as long as *k* is large enough and *v* is not too large, this condition can be satisfied.

**Success-biased social learning model**. The success-bias model uses the same abstract formalism for preferential transmission of adaptive behavior as for parasitic behavior. Specifically, assume that cues of success are available that are specific to the adaptive component of a variant. This means that whether or not a variant is also parasitic does not influence a cue of its success at achieving some task. These cues may be material evidence, such as larger crop yields or longer-lasting clothing or healthier children, or merely verbal testimony. Let *σ* be the proportional increase in odds of acquiring behavior that was adaptive in the *previous* generation. Simultaneously and independently, the same system of preferential transmission of parasitic behavior applies, and so *w* is again the proportion odds of acquiring parasitic behavior. This leads to the following modifications to the probabilities of acquiring each variant combination by social learning:

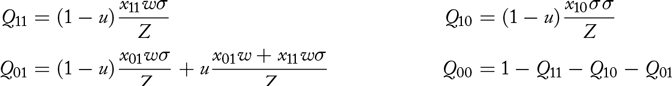
where *Z* = *x*_11_*w σ* + *x*_10_ *σ* + *x*_01_*w* + *x*_00_ is a common normalizing factor. The fitness of this success-biased social learning strategy is:

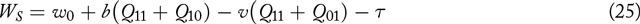
where τ is the cost of searching for and using the success cues.

Let *p* be the frequency of the success-bias allele, with the remainder 1 − *p* assigned for the moment only to pure individual learning. The recursions for *q*, *r*, and the covariance *k* are:

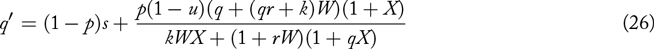

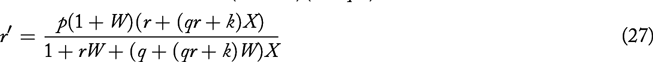

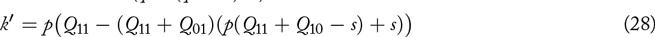
where *W* = *w* − 1 and *x* = a − 1 are substituted for the sake of compact notation.

There are four sets of solutions for *q*, *r*, and *k* for which the expected values remain unchanged. For two of these solutions, there is a non-zero covariance at steady state. For the first of these solutions, 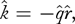 as in the analogous solution for the critical social learning model. For the final solution, the expected steady state covariance is given by:

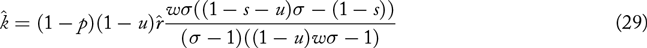

Note the leading term 1 − *p*. In the absence of pure individual learners, the covariance goes to zero. This is because the expected value of parasitic behavior goes to 1 in that event, and so there is no variance and so no covariance. But as long as innovation maintains variation at steady state, this expression can be positive, provided both

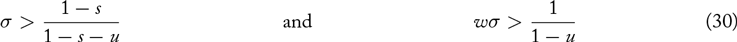

The intuition behind the positive covariance in this model is that the product *wσ* favors combinations of adaptive and parasitic behavior. This combination is transmitted at a higher rate than any other. This builds covariance. So the mechanism is quite different than the filtering mechanism from the critical social learning model. But the result is analogous.

A curious obstacle in interpreting this model is that success-bias, like conformist transmission, can actually exclude all innovation from the population. More commonly, it can reduce the innovation rate to negligible levels. While this result is instructive—strategic social learning can substitute for innovation—it is not easily interpreted as a biologically realistic result. Some force of innovation, whether deliberate or by mistake, seems to be required.

There are many ways to define success-biased social learning. It would be possible to instead allow success-bias to use a proxy of total fitness of a social model, and therefore incorporate the parasite cost *v* into its cue. However doing so would sterilize by assumption any fertile hypothesis invoking parasitic behavior. A more interesting idea perhaps is to consider that beliefs about the payoff value of behavior are themselves socially learned, especially when payoffs are distantly removed from the actions that influence them.

## References

Boyd, R. and Richerson, P.J. (1985). Culture and the Evolutionary Process. University of Chicago Press.

Boyd, R. and Richerson, P. J. (1995). Why does culture increase adaptability? Ethology & Sociobiology, 16:125–143.

Boyd, R. and Richerson, P. J. (1996). Why culture is common, but cultural evolution is rare. Proceedings of the British Academy, 88:77–83.

Boyd, R. and Richerson, P. J. (2000). Meme theory oversimplifies how culture changes. Scientific American, 283: 70–71.

Cavalli-Sforza, L. L. and Feldman, M. W. (1981). Cultural transmission and evo-lution: a quantitative approach. Princeton University Press.

Dawkins, R. (1976). The Selfish Gene. Oxford University Press.

Durham, W. H. (1991). Coevolution: Genes, culture, and human diversity. Stanford University Press.

Enquist, M., Eriksson, K., and Ghirlanda, S. (2007). Critical social learning: A solution to Rogers’s paradox of nonadaptive culture. American Anthropologist, 109:727–734.

Healey, K. (2016). Fuck nuance. Sociological Theory.

Henrich, J. and McElreath, R. (2003). The evolution of cultural evolution. Evolutionary Anthropology, 12:123–135.

Kendal, J. R. and Laland, K. N. (2000). Mathematical models for memetics. Journal of Memetics, 4.

Laland, K. N. (2004). Social learning strategies. Learning and Behavior, 32:4–14.

Lloyd-Smith, J., Schreiber, S. J., Kopp, P. E., and Getz, W. M. (2005). Superspreading and the impact of individual variation on disease emergence. Nature, 540 pages 355–359.

McElreath, R. (2012). The coevolution of social learning and sensitivity to changing environments.

McElreath, R. and Strimling, P. (2008). When natural selection favors learning from parents. Current Anthropology, 49:307–316.

McElreath, R., Wallin, A., and Fasolo, B. (2013). The evolutionary rationality of social learning. In Hertwig, R., Hoffrage, U., and the ABC Research Group, editors, Simple Heuristics in a Social World, pages 381–408. Oxford University Press.

Nakahashi, W., Wakano, J. Y., and Henrich, J. (2012). Adaptive social learn-ing strategies in temporally and spatially varying environments. Human Nature, 23:386–418.

Richerson, P. J. and Boyd, R. (2005). Not by genes alone: How culture transformed human evolution. University of Chicago Press.

Rogers, A. R. (1988). Does biology constrain culture? American Anthropologist, 90:819–831.

